# Intralocus sexual conflict can maintain alternative reproductive tactics

**DOI:** 10.1101/2021.11.26.470147

**Authors:** Madilyn Marisa Gamble, Ryan G Calsbeek

## Abstract

Alternative reproductive tactics (ARTs) are ubiquitous throughout the animal kingdom. Multiple mechanisms have been proposed to explain the maintenance of ARTs over time, including disruptive and negative frequency-dependent selection and female choice. However, these mechanisms focus exclusively on selection in the sex exhibiting the polymorphism, potentially limiting our understanding of the evolutionary causes and consequences of ARTs. Here we explore the role that intralocus sexual conflict may play in the maintenance of sex-specific ARTs. We use a genetically explicit individual-based model in which female fecundity and male tactic share a genetic architecture. By modeling ART maintenance under varying selection regimes and levels of sex-specific gene expression, we explore the conditions under which intralocus sexual conflict can maintain a hypothetical ART defined by two color morphs that depend on an underlying liability trait. Our models consistently revealed that sexual conflict can result in the persistence of a sex-specific polymorphism over hundreds of generations, even in the absence of negative frequency-dependent selection. ARTs were maintained through correlated selection when one male ART has lower fitness but produces daughters with higher fitness. Importantly, the maintenance of ARTs through sexual conflict resulted in a significant reduction in population growth rate, indicating that the evolutionary mechanism of ART maintenance can have broad ecological consequences. These results highlight the importance of understanding selection on both sexes when attempting to explain the maintenance of ARTs. Our results are consistent with a growing literature documenting genetic correlations between male ARTs and female fitness, suggesting that the maintenance of sex-specific ARTs through intralocus sexual conflict may be common and widespread in nature.

## Introduction

Alternative reproductive tactics (ARTs) are ubiquitous throughout the animal kingdom and their evolution is favored by high variance in reproductive success (Shuster and Wade 2003). Proximate mechanisms underlying ARTs include purely genetically based polymorphisms (Shuster 1989; Cordero 1990; Lank et al. 1995; Sinervo and Lively 1996; Sinervo et al. 2000), purely environmentally induced polymorphisms (Thornhill 1981; Emlen 1994; Hunt and Simmons 1997; Moczek and Emlen 1999), as well as those mediated by a combination of genetic and environmental factors (Neff and Svensson 2013). Multiple mechanisms have been proposed to explain the evolutionary maintenance of ARTs over time, including disruptive selection, negative frequency-dependent selection, and female choice (Wright 1969; Clarke and O’Donald 1974; Gross 1985, 1991, 1996; Fitzpatrick et al. 2007; Hughes et al. 2013); for review, see (Ayala and Campbell 1974; Oliveira et al. 2008). However, these forms of selection focus only on the sex expressing the polymorphism – often males –ignoring the fact that females may also carry the genes underlying male-specific ARTs (Sinervo and Zamudio 2001; Barson et al. 2015; Heinen-Kay et al. 2019a; Pearse et al. 2019). Theory and empirical studies have shown that interlocus sexual conflict affects the evolution and maintenance of ARTs (Alonzo and Warner 1999, 2000; Alonzo 2007), but the potential role of intralocus sexual conflict has yet to be considered. By ignoring how selection operates on females carrying the genetic architecture of male ARTs, we may limit our understanding of the diversity of mechanisms maintaining sex-specific polymorphisms, as well as their evolutionary causes and consequences.

Indeed, a growing body of evidence shows that selection on male ARTs and the traits that differentiate them results in correlated evolution in females: genetic linkage between male ARTs and female reproductive traits and fitness has been described in lizards (Sinervo and Zamudio 2001), crickets (HeinenLJKay et al. 2020; Richardson et al. 2021), and bulb mites (Bielak et al. 2014; Buzatto et al. 2018; Buzatto and Clark 2020; Łukasiewicz et al. 2020). This genetic linkage, coupled with sexually antagonistic selection (Lande 1980), leads to intralocus sexual conflict (Bonduriansky and Chenoweth 2009), in which the genes underlying one male tactic lead to a decrease in fitness when carried by females. While others have proposed that intralocus sexual conflict may maintain additive genetic variance (Rice and Chippindale 2001; Foerster et al. 2007) and genetic polymorphism (Connallon and Clark 2012), result in balancing selection (Connallon and Clark 2014), and promote speciation (Bonduriansky 2011), the potential for intralocus sexual conflict to maintain alternative reproductive tactics is comparatively understudied. Nevertheless, the variety of systems in which intralocus sexual conflict occurs between certain male ARTs and female reproductive traits suggests that intralocus sexual conflict could play a role in ART maintenance, especially because both unresolved intralocus sexual conflict and ARTs are common in nature (Oliveira et al. 2008; Cox and Calsbeek 2009).

While existing analytical models describe the action of selection on quantitative traits (Otto and Day 2007), correlated traits (Lande and Arnold 1983), and threshold traits including ARTs (Gross and Repka 1998), we are unaware of any theory that simultaneously investigates evolution under both sexually antagonistic selection on quantitative threshold traits (which often underly ARTs; (Roff 1996) and variable intersexual genetic correlations (which control the degree of sexual conflict; (Rice and Chippindale 2001; Bonduriansky and Chenoweth 2009). Here we introduce a proof of concept model (Servedio et al. 2014) that takes new steps towards understanding the role that intralocus sexual conflict may play in the maintenance of sex-specific ARTs. We use a genetically explicit individual-based model in which a quantitative liability trait influences both female fecundity and male tactic through a shared genetic architecture. By modeling ART maintenance under varying selection regimes and levels of sex-specific gene expression, we explore the conditions under which intralocus sexual conflict can maintain a hypothetical male ART defined by two color morphs (purple and yellow). In this theoretical model, the genes for purple males confer high fecundity to females while the genes for yellow males confer low fecundity to females.

To evaluate the potential for sexually antagonistic selection and intralocus sexual conflict to maintain ARTs, we allowed a theoretical population to evolve under a range of scenarios representing three different selection regimes acting on males: no selection, directional selection against purple males, and negative frequency-dependent selection. We included negative frequency-dependent selection both to show that the model correctly captures processes we know can maintain ARTs, and to compare the outcome of negative frequency-dependent selection to the outcome of intralocus sexual conflict. Within each selection regime we also varied the degree to which males and females could evolve independently by varying the number of genetic loci with sex-dependent expression. We used these models to test our hypothesis that intralocus sexual conflict may influence ART frequency over time, and that this process might be altered by different selection regimes and/or the breakdown of intersexual genetic correlations through sex-specific gene expression. Specifically, we predicted ARTs could be maintained: (1) when purple and yellow males have equal fitness but intersexual heritability is low such that more fecund females do not necessarily produce purple sons, and (2) when purple males have lower fitness than yellow males but intersexual heritability of the liability trait is high such that low-fitness purple males produce highly fecund daughters. We also investigated how the maintenance of ARTs through intralocus sexual conflict could affect population growth rates through the maintenance of the sex load, which occurs when neither sex can reach its phenotypic optimum (Rice and Chippindale 2002). Because a sex load decreases the fitness of each sex, we predicted that populations growth rates would decrease when ARTs were maintained through sexual conflict.

## Methods

### Overview

In modeling the frequency of two different male ARTs – defined by purple and yellow male color morphs – we assumed that ART was defined by a quantitative threshold trait, which in turn was entirely genetically controlled by 20 quantitative loci. Individual males whose liability trait value was greater than or equal to a threshold value became purple males, and those below the threshold became yellow males. Females, which did not express these ARTs, had a fecundity value proportional to the same underlying quantitative trait. For simplicity we assumed a discrete population of semelparous individuals with non-overlapping generations. We did not include linkage disequilibrium or mutations in our model; all genetic variance stemmed from an initial random draw of allele values in the first generation.

To test our hypothesis that intralocus sexual conflict can affect the maintenance of ARTs, we varied both the form of selection acting on the male ARTs and the intersexual heritability for the quantitative liability trait (hereafter “liability trait”). We varied the form of selection on males by changing the relative fitness of each tactic. We exposed the population to a null selection regime in which both male ARTs were equally likely to survive and mate. We simulated sexually antagonistic selection by assigning lower fitness to purple males but higher fitness (fecundity) to females with higher liability trait values. Finally, we modeled negative frequency-dependent selection in which the rare male ART in each generation had a fitness advantage to compare this established method of ART maintenance to the new mechanism proposed here.

We altered the intersexual heritability of the liability trait by varying the number of loci that were “general” or “sex-specific” in their expression. When all 20 loci were “general,” the effect of each allele controlling the liability trait was the same in both sexes. This made the liability trait equally heritable within and between the sexes, so purple males produced more fecund daughters than yellow males. We predicted that correlated selection acting on males because of fecundity selection might allow purple males to persist in the population even when directional selection acts against them. However, as more loci become “sex-specific” – that is, allelic expression depended on sex and genetic architecture for the liability trait was less shared between the sexes – intersexual heritability will decline, and sire tactic should no longer influence daughter fitness. Thus, purple males should no longer necessarily produce fitter daughters, and we predicted that consistent directional selection against purple males could result in their decline or disappearance from the population.

### The Model

We initiated each simulation with a parent generation comprised of 1000 diploid sexually reproducing individuals (50:50 sex ratio). Each individual in our model had two traits that made up its phenotype: sex and the liability trait. Sex was assigned randomly, and the liability trait was a quantitative trait determined by the sum of allele values measured across 20 loci. To allow for sex-specific allelic effects, each allele had two possible values (*f* and *m*) that influenced the liability trait. That is, each locus had a total of three possible values that contribute to the liability trait, and which value was expressed depends on whether the locus was sex-specific or general and if the former, whether the individual was male or female. For each sex-specific locus, only the two allele values corresponding to that individual’s assigned sex contributed to the liability trait value. This is the biological equivalent of an allele that has sex-specific expression, for example, through sex hormone regulation (Rice and Chippindale 2001). For general loci the value of each allele at that locus was determined by averaging the allele’s two possible values. Allele values for the initial generation were drawn randomly from a normal distribution with mean = 0 and SD = 0.5. We calculated the liability trait value (*Y*) of the individual from its genotype using the following sex-specific equations:

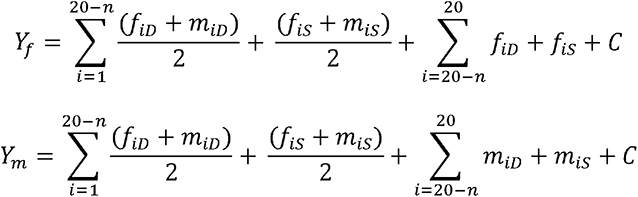

where *n* was the number of sex-specific loci, *f* and *m* are the female- and male-specific values, respectively, of the two alleles at the *i*th locus inherited from the dam (*D* subscript) or the sire (*S* subscript), and *C* is a constant. The first summation calculated the component of the liability trait value contributed from the general loci; the second summation calculated the contribution of the sex-specific loci. Adding a constant to the expression ensured that the liability trait value remained a positive number with a mean of *C* in the parent generation; any individual whose calculated the liability trait value was less than zero was assigned a value of zero. We arbitrarily set *C* = 10 for our simulations.

Male mating tactic was determined by threshold liability trait value: males with a value greater than or equal to 10 were purple males and those with a value less than 10 were yellow males. Fecundity (*E*) was assigned to females as a function of the liability trait value:

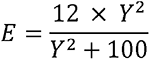

such that fecundity increased asymptotically with increasing liability trait value to a maximum value approaching 12 offspring at a liability trait value of 40. This set an upper limit on fecundity selection while preserving positive directional selection on the liability trait in females.

We modeled the evolution of male and female liability trait values and ART frequency under seven different selection regimes on male tactic and five different values for the number of sex-specific loci for a total of 35 different model scenarios (Table 1). These varied in the degree to which selection on the liability trait was sexually antagonistic and in the intersexual heritability of the liability trait, thus creating a spectrum of intralocus sexual conflict. We varied direct selection on males by changing the relative combined probability of survival and mating (hereafter *W_T_* for each tactic). In the “no selection” model individuals of both ARTs had equal values of *W*. In three separate “directional selection” models we set *W_p_* = 1 and *W_y_* = 1.5, 1.8, or 2. This allowed us to evaluate how the strength of direct selection on male tactic affected ART maintenance in the face of intralocus sexual conflict. In each directional selection model, *W_p_* and *W_y_* were held constant across generations. In three separate “negative frequency-dependent selection” models, each tactic’s *W* varied as function of its frequency (*F_T_*) such that the rare tactic in each generation always had a higher mating probability (1- *F_T_*). We ran three different negative frequency-dependent models that differed in starting frequency ratios of purple and yellow males: *F_p_:F_y_ =* 2:1, 1:1, and 1:2. This allowed us to evaluate how initial tactic frequencies affected ART maintenance under negative frequency-dependent selection.

**Table 1.** Parameters and values modeled, resulting in 35 total scenarios modeled. Each scenario varied in the form of selection on males and the number of sex-specific loci. Negative frequency- dependent models also varied in the starting frequency ratio of purple to yellow males (*F_p_:F_y_*); directional selection models also varied in the ratio of purple fitness to yellow fitness (*W_p_:W_y_*).

| Parameter | Values modeled |
| --- | --- |
| Male Selection | No Selection, $W_p = W_y$ |
| Regime | Negative frequency-dependent selection, $F_p:F_y = 1:1$ |
| | Negative frequency-dependent selection, $F_p:F_y = 1:2$ |
| | Negative frequency-dependent selection, $F_p:F_y = 2:1$ |
| | Directional selection against purple males, $W_p:W_y = 1:1.5$ |
| | Directional selection against purple males, $W_p:W_y = 1:1.8$ |
| | Directional selection against purple males, $W_p:W_y = 1:2$ |
| # Sex-Specific Loci | 0, 5, 10, 15, 20 |

In all models, *W_f_*did not differ among individual females, but the proportion of females that survived to mate in each generation declined with population size to simulate density-dependent population growth regulation. The proportion of females surviving to mate (*M_f_*) was calculated anew for each generation using the equation:

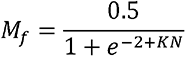

where *K* is a constant and *N* is the current population size. We chose *K* = 0.0015 such that this equation defined a sigmoidal curve that decreased from 0.44 at *N* = 0 to 0.04 at *N* = 3000. For each generation the number of females that were allowed to reproduce (*D*) was calculated as:

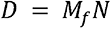

*D* females were randomly chosen to be dams. *D* males were similarly chosen to be sires but sire selection was weighted by *W_T_*. Dams were randomly paired with sires (no assortative mating), and a number of offspring equivalent to the dam’s fecundity were created. Each offspring inherited alleles at all 20 loci in a Mendelian fashion such that at each locus the offspring inherits one allele from the dam and one from the sire, and the allele inherited from each parent is randomly drawn from the two alleles each parent possesses. Sex was assigned randomly to offspring to achieve an approximately equal sex ratio, and the liability trait value, fecundity, and mating probability were calculated for each offspring as described above for individuals in the parent generation. These offspring were then used as parents for the next generation, and the model was iterated for a total of 500 generations in each model scenario. We ran 11 replicates of the full set of the 35 model scenarios to ensure that results did not qualitatively differ among model runs.

### Analysis

In each of the 35 model scenarios, for each generation we calculated (1) intra- and intersexual heritability of the liability trait value (to understand the degree to which males and females could respond independently to selection), (2) the sex-specific mean and variance of the liability trait value (to understand the degree of sexual dimorphism that evolved and the opportunity for selection), (3) sex-specific selection gradients for the liability trait (to understand the degree to which selection was sexually antagonistic), and (4) the frequency of ARTs (to understand how intralocus sexual conflict affects ART maintenance). Intra- and intersexual heritabilities were calculated by doubling the regression coefficient of the sex-specific mean offspring liability trait value regressed on sex-specific parent liability trait value (Falconer 1989). Sex-specific selection gradients for each generation were calculated as the slope of the regression of relative fitness on standardized liability trait value. Relative fitness for each sex was calculated by dividing the number of offspring produced by each individual by the mean number of offspring produced by that sex; standardized liability trait value was calculated by scaling each individual’s liability trait value to the sex-specific mean and dividing by the sex-specific standard deviation. We then recorded how each of these factors changed across generations (i.e., as additive genetic and phenotypic variance changed), and how that change varied with (1) the selection regime and (2) the number of loci that were sex-specific rather than general in their effect on the liability trait. Finally, intralocus sexual conflict is known to affect intrinsic population growth rates through the maintenance of the sex load (Rice and Chippindale 2002). To understand how population growth rates are affected by the maintenance of ARTs through intralocus sexual conflict, we also plotted population size over 500 generations for each selection regime and calculated geometric mean population growth rates (Lewontin and Cohen 1969) for each selection regime and number of sex-specific loci.

## Results

Results were not qualitatively different among the 11 replicate sets of model runs. Therefore we report results from a single haphazardly chosen set of 35 model runs.

Increasing the number of sex-specific loci decreased intersexual heritability. Intrasexual heritability was approximately 1 and was constant across selection regimes, numbers of sex-specific loci, and generations. Intersexual heritability generally decreased from 1 to 0 as the number of sex-specific loci increased from 0 to 20 (Figure 1). This trend was consistent across all selection regimes.

**Figure 1.**
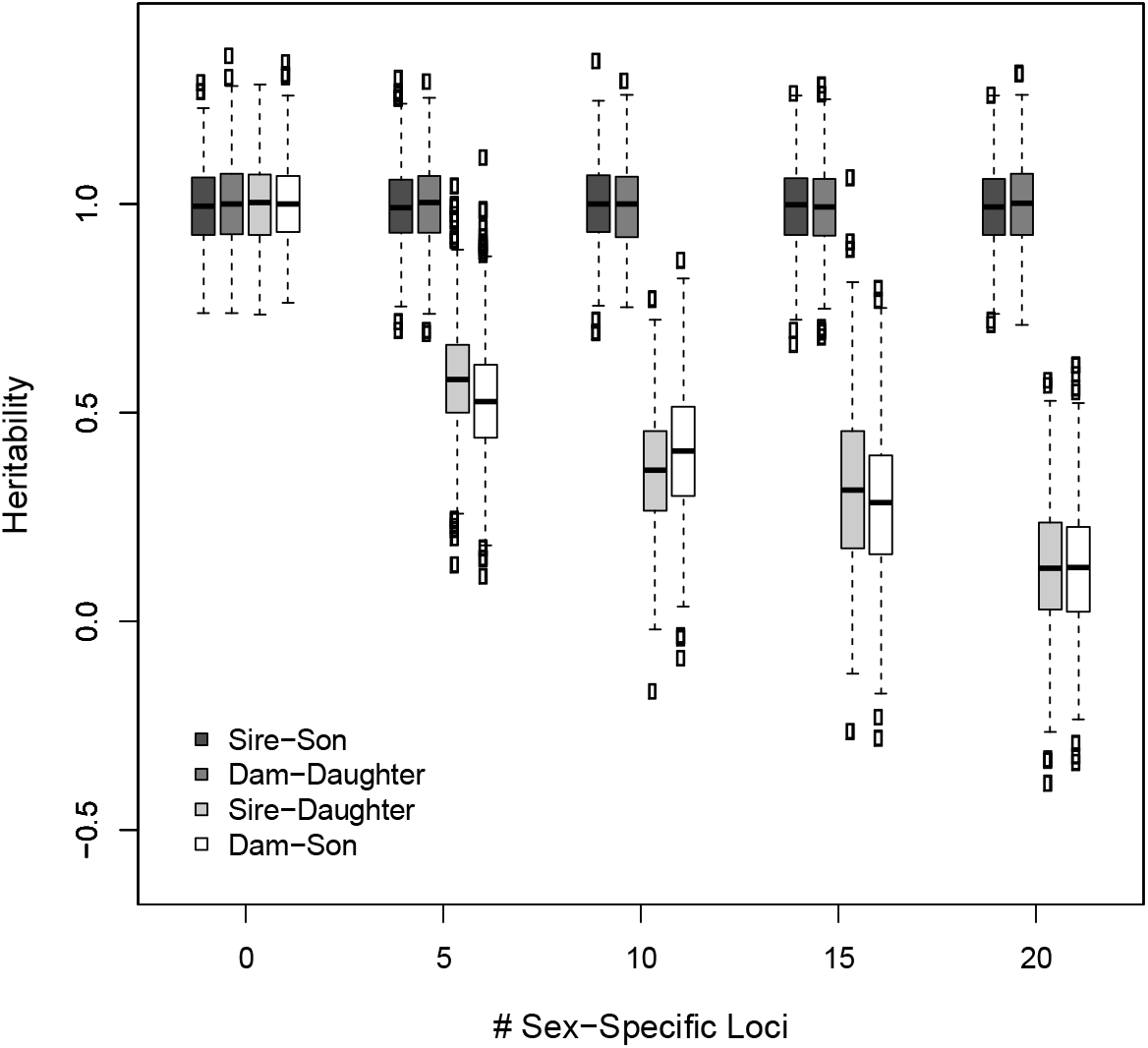
Box plots show mean values of inter- and intrasexual heritability averaged across 500 generations in a “no selection” simulation. Intersexual heritability declined as the number of sex-specific loci increased. Intrasexual heritability for the liability trait value did not vary with the number of sex-specific loci. Results were qualitatively similar across all other selection regimes. Solid line indicates median, box encompasses the interquartile range, dashed lines indicate minimum and maximum values, and circles indicate outliers.

Increasing the number of sex-specific loci also facilitated the evolution of sexual dimorphism in the liability trait value. Sexual dimorphism evolved in all selection regimes when intersexual heritability was less than 1 (>0 sex-specific loci; Figure 2). The rate and degree to which the sexes diverged in their liability trait value depended on the selection regime and the number of sex-specific loci (i.e., the intersexual heritability). In all cases, sexual dimorphism evolved faster and to a greater degree when intersexual heritability was low (more sex-specific loci) and when selection was sexually antagonistic (males experienced directional selection favoring yellow males; Figure 2c). This result is consistent with existing literature and theory on the evolution of sexual dimorphism (Cox and Calsbeek 2009).

**Figure 2.**
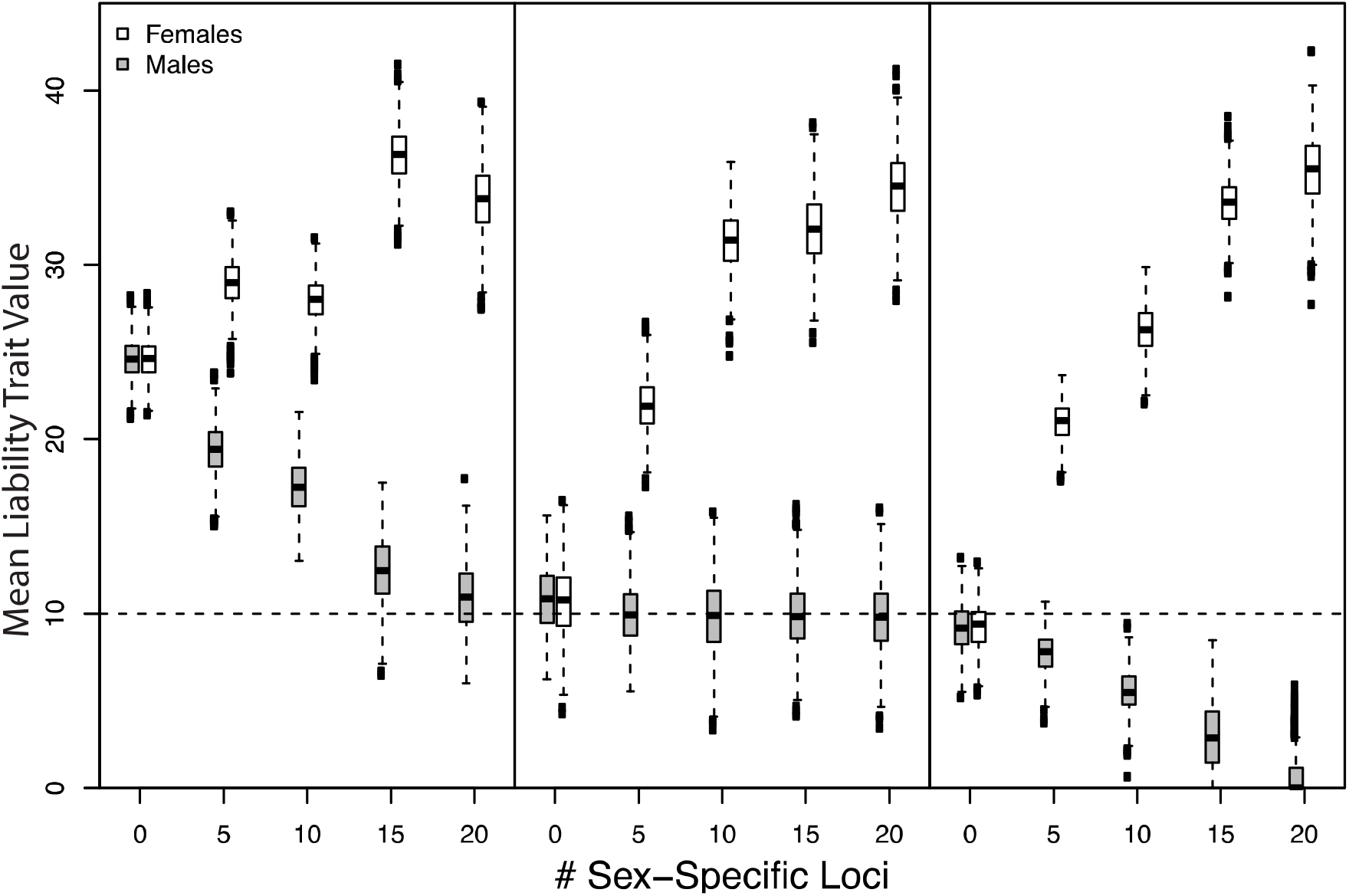
Box plots showing the final mean liability trait value of females and males across a range of the number of sex-specific loci in (a) a no selection regime, (b) a negative frequency-dependent selection regime in which initial tactic frequencies were equal, and (c) a directional selection regime in which yellow males were 1.8 x more likely to survive and mate than purple males. Dotted horizontal line represents the size threshold separating purple and yellow males, therefore male boxplots overlapping this line indicate that both tactics persisted over 500 generations in that scenario. Results from negative frequency-dependent selection regimes with different initial frequencies and from directional selection regimes with different mating probability ratios are qualitatively the same as (b) and (c), respectively, and are not shown here. Solid line indicates median, box encompasses the interquartile range, dashed lines indicate minimum and maximum values, and circles indicate outliers.

Sex-specific selection gradients reflected the imposed selection regimes initially in all selection regimes (Figure 3). In later generations, selection gradients for both males and females converged on zero in most models, as the population responded to selection and the liability trait evolved. Shallower selection gradients in later generations reflected a reduction in the variance in male liability trait values (as the trait responded to selection), a reduction in the covariance between female liability trait value and fecundity (as females reach a trait value at which the fecundity-trait value relationship asymptotes), or both. Differences in sex-specific selection gradients were maintained in the directional and negative frequency-dependent selection regimes in which intersexual heritability was high (zero sex-specific loci) and ARTs were maintained (see below).

**Figure 3.**
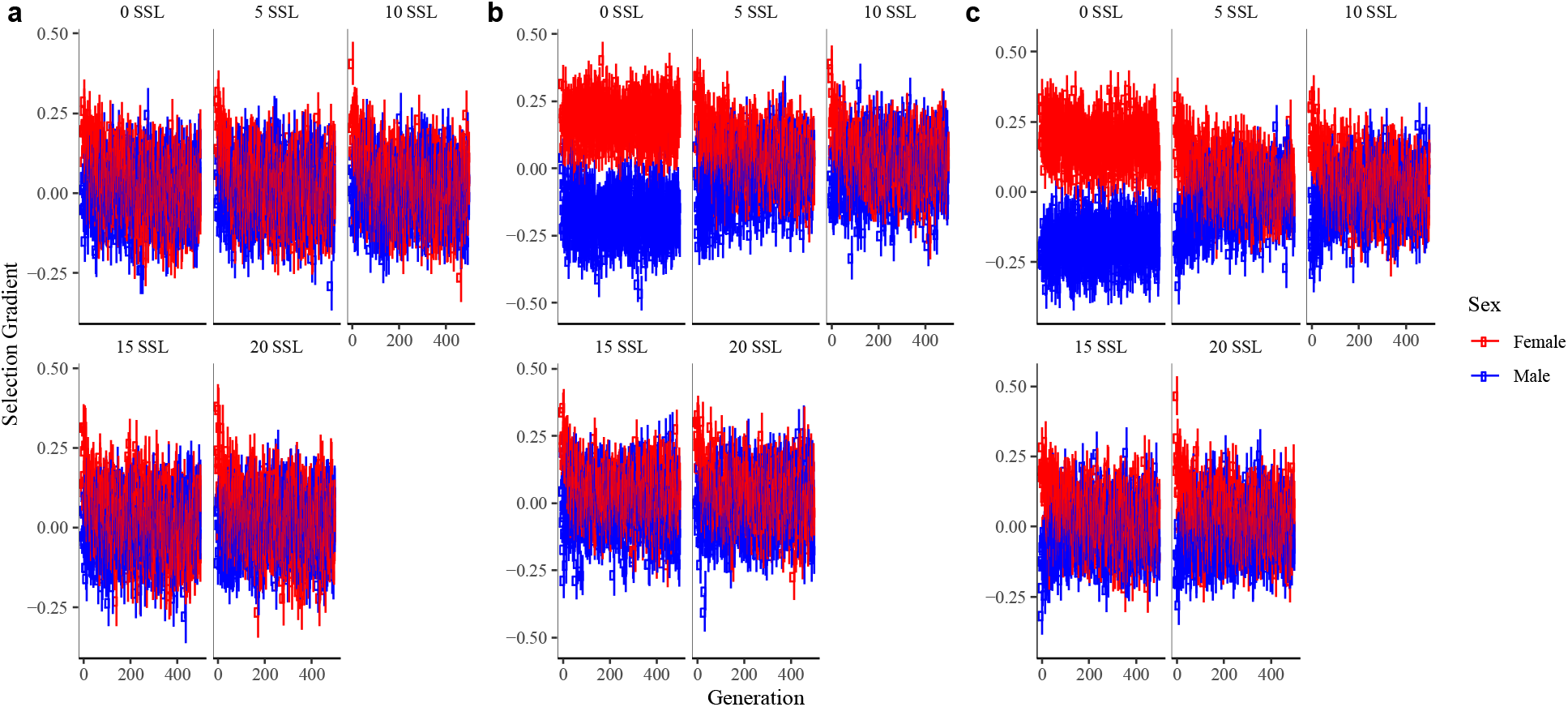
Selection gradients on the liability trait in males and females over 500 generations and across a range of the number of sex-specific loci (SSL) in (a) a no selection regime, (b) a negative frequency-dependent selection regime in which initial tactic frequencies were equal (*F_p_*=*F_\y_*), and (c) a directional selection regime in which yellow males were 1.8 x more likely to survive and mate than purple males. Differences in sex-specific selection gradients were maintained in (b) the negative frequency-dependent and (c) directional selection regimes in which intersexual heritability is high (zero sex-specific loci), constraining the evolution of sexual dimorphism and maintaining the sex load.

Selection regimes and intersexual heritabilities affected geometric mean population growth rates. In most models, population size increased from 1000 in the parent generation to an asymptote of approximately 2000 individuals (Figure 4). However, in all the directional selection and negative frequency-dependent selection regimes, population size and geometric mean population growth rate were greatly reduced when there were zero sex-specific loci (i.e., when intersexual heritabilities were approximately 1). In no-selection regimes, geometric mean population growth rates did not differ across numbers of sex-specific loci. In negative frequency- dependent and directional selection regimes, geometric mean population growth rates differed across numbers of sex-specific loci such that they were lowest when the number of sex-specific loci was zero. The degree to which population growth rate was reduced when there were no sex-specific loci (compared to other values of sex-specific loci) did not vary with different starting frequencies in the negative frequency-dependent selection models. However, in directional selection regimes with zero sex-specific loci (i.e. models with intralocus sexual conflict), population growth rates decreased as the strength of directional selection against purple males increased. These reductions in population size and geometric mean population growth rate were consistent with the unresolved sex load maintained when males and females were not able to evolve independently (Arnqvist and Tuda 2010). The maintenance of the sex load over time was evident in the sex-specific selection gradients for the directional selection simulations with no sex-specific loci (Figure 3).

**Figure 4.**
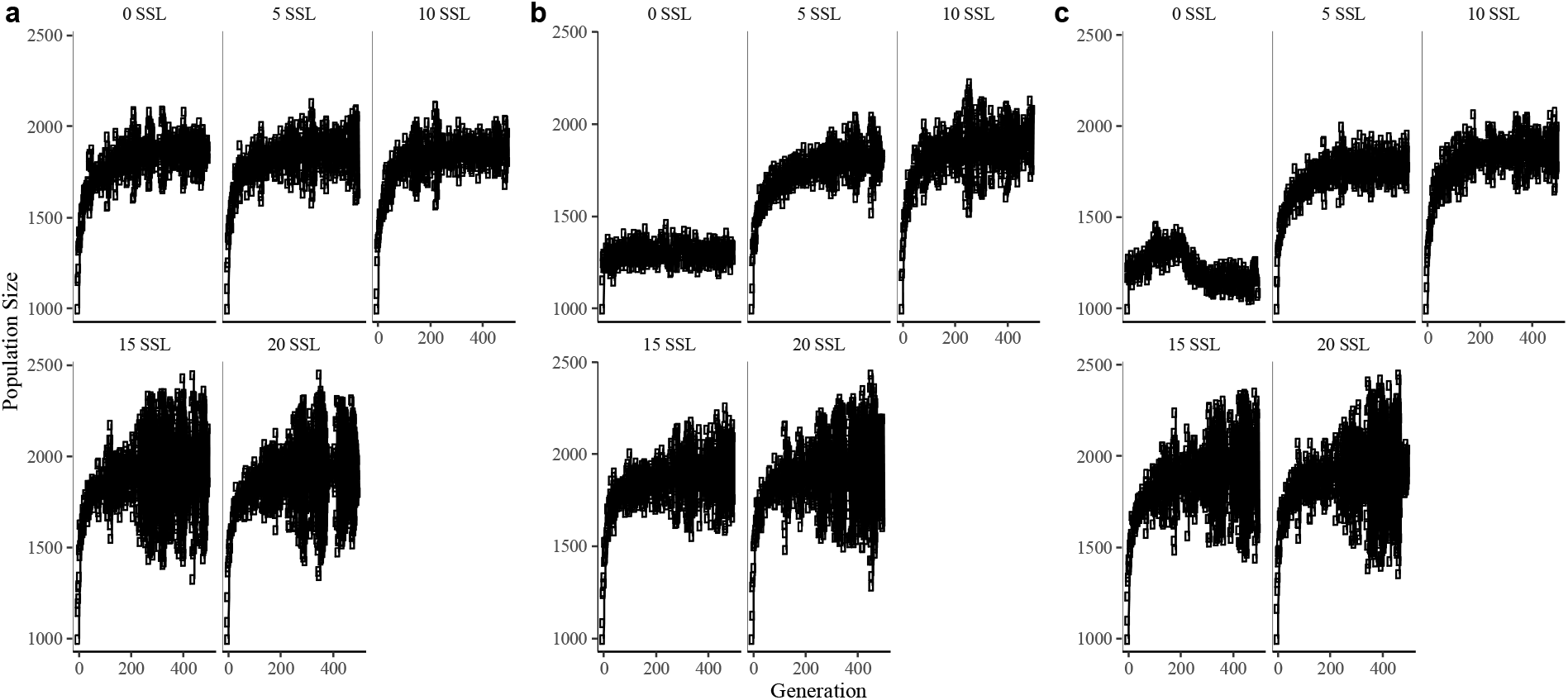
Population sizes over 500 generations and across a range of the number of sex-specific loci (SSL) in (a) a no selection regime, (b) a negative frequency-dependent selection regime in which initial tactic frequencies were equal (*F_p_*=*F_y_*), and (c) a directional selection regime in which yellow males were 1.8 x more likely to survive and mate than purple males. In (b) the negative frequency-dependent and (c) directional selection regimes in which intersexual heritability is high (zero sex-specific loci), population size is reduced because of the sex load.

Alternative reproductive tactics were maintained under three different conditions. First, under negative frequency-dependent selection (i.e., the probability of mating for tactic *T* is 1-*F_T_*), both tactics were maintained regardless of initial tactic frequencies or the degree of intersexual heritability (Figure 5). This result was consistent across all 11 replicate model runs.

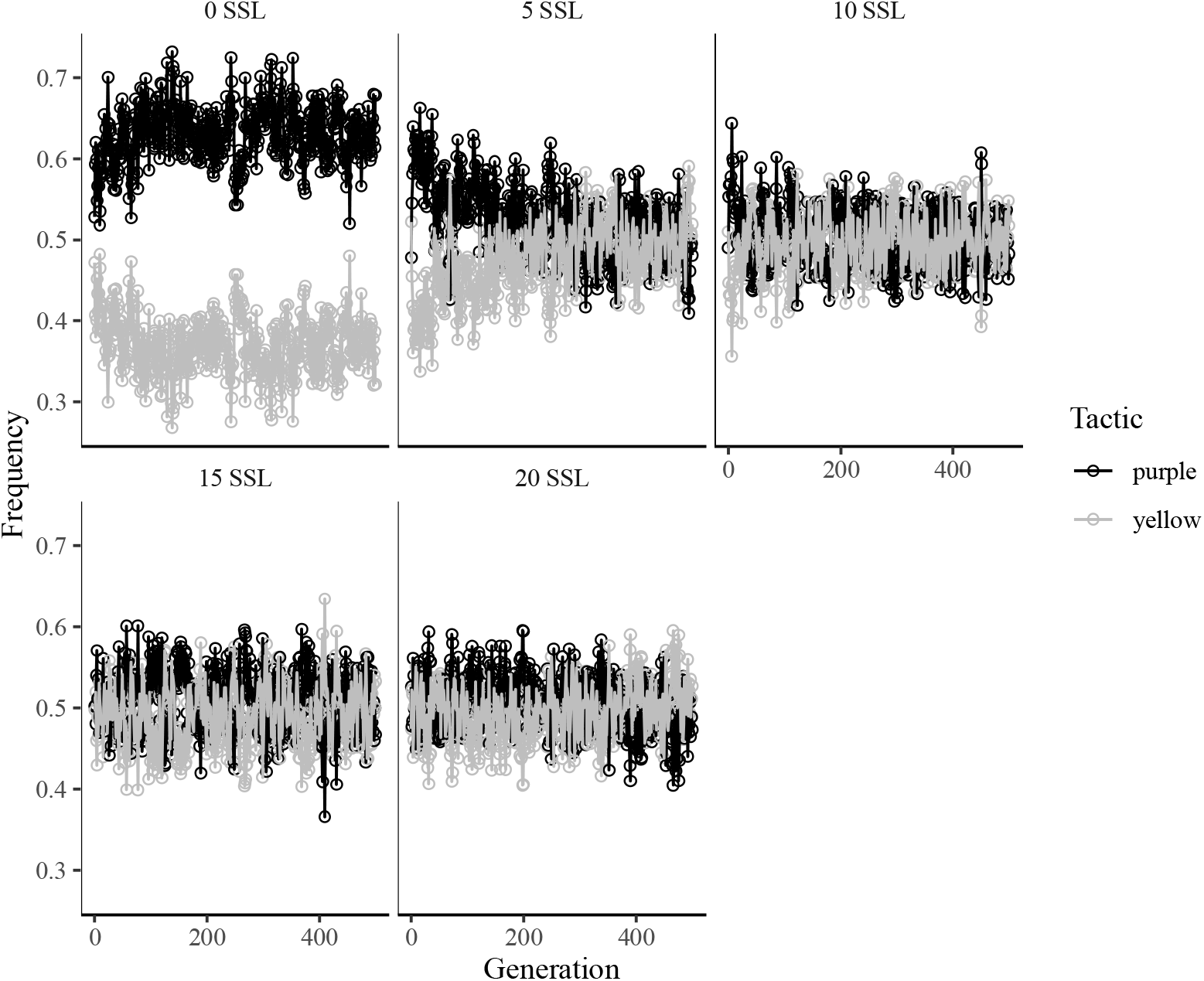

There were two general situations besides negative frequency-dependent selection in which ARTs were preserved across 500 generations in our model. First, ARTs were maintained in the no selection regime when intersexual heritability was low (e.g., 15 or 20 sex-specific loci; Figure 6). Under no selection and 15 sex-specific loci, both tactics persisted over 500 generations in 9 of 11 model runs; under no selection and 20 sex-specific loci, both tactics persisted over 500 generations in 10 of 11 model runs. In these same models, purple males comprised >75% of the population by the 500^th^ generation in 10 out of 11 “no selection” model runs with 15 sex-specific loci and 5 out of 11 “no selection” model runs with 20 sex-specific loci. Across all 11 “no selection” model run replicates, increasing the number of sex-specific loci resulted in a slower trend toward fixation of purple males (Figure 6).

**Figure 6.**
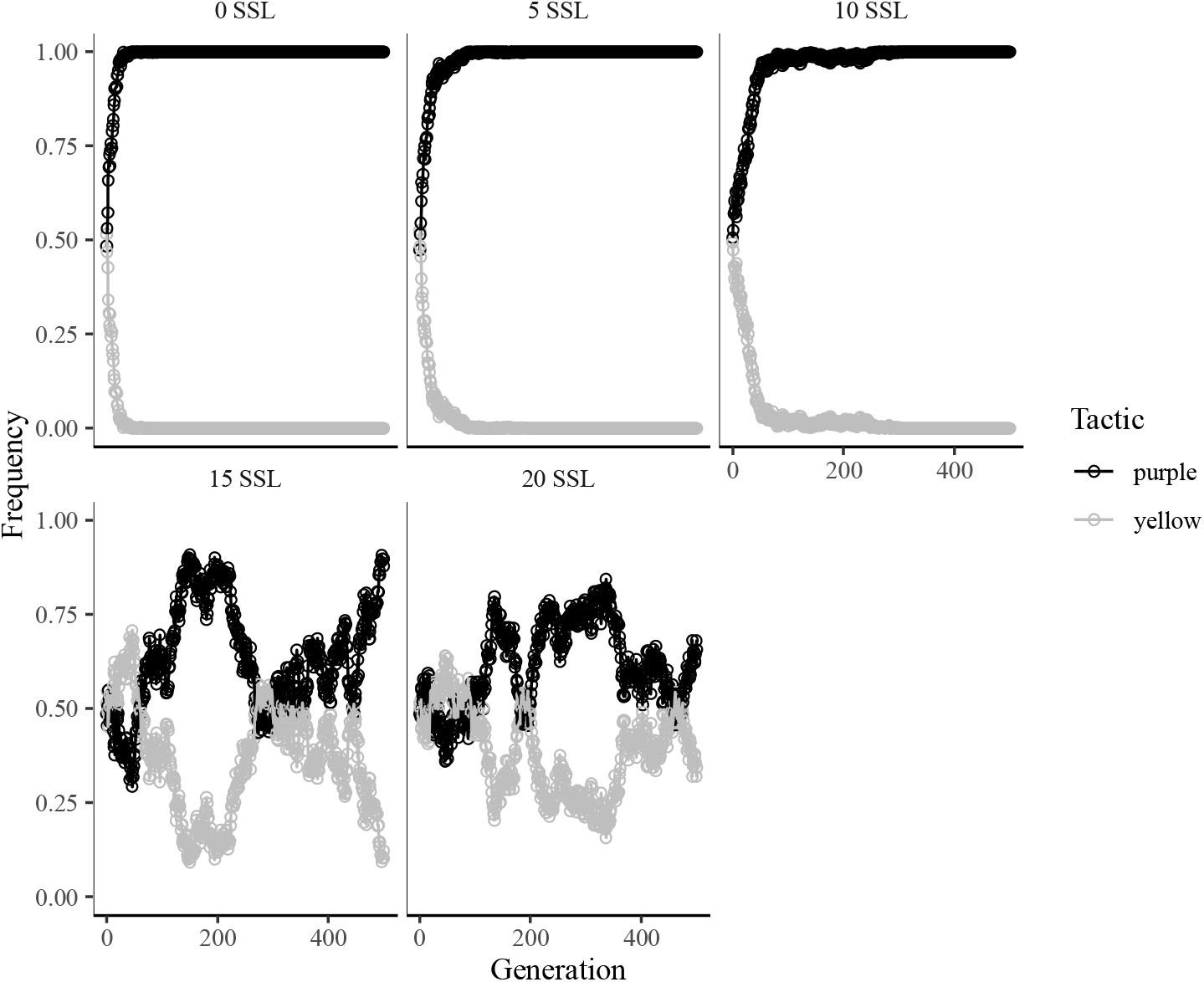
Frequency of purple and yellow males over 500 generations and across a range of the number of sex-specific loci (SSL) in a “no selection” regime.

ARTs were also maintained by intralocus sexual conflict over the liability trait. This occurred when selection on the liability trait was sexually antagonistic (favoring males with a small liability trait value and females with a large liability trait value) and intersexual heritability was high (e.g., zero sex-specific loci / all loci affect males and females equally; Figure 7). This was especially evident under strong directional selection (e.g., when yellow males were 1.8 – 2 times more likely to survive and reproduce than purple males. In this situation ARTs were maintained by the balance between (a) fecundity selection favoring large liability trait value in females and high intersexual heritability producing purple sons and (b) selection against those purple sons. This outcome was consistent across all 11 replicates of our model runs.

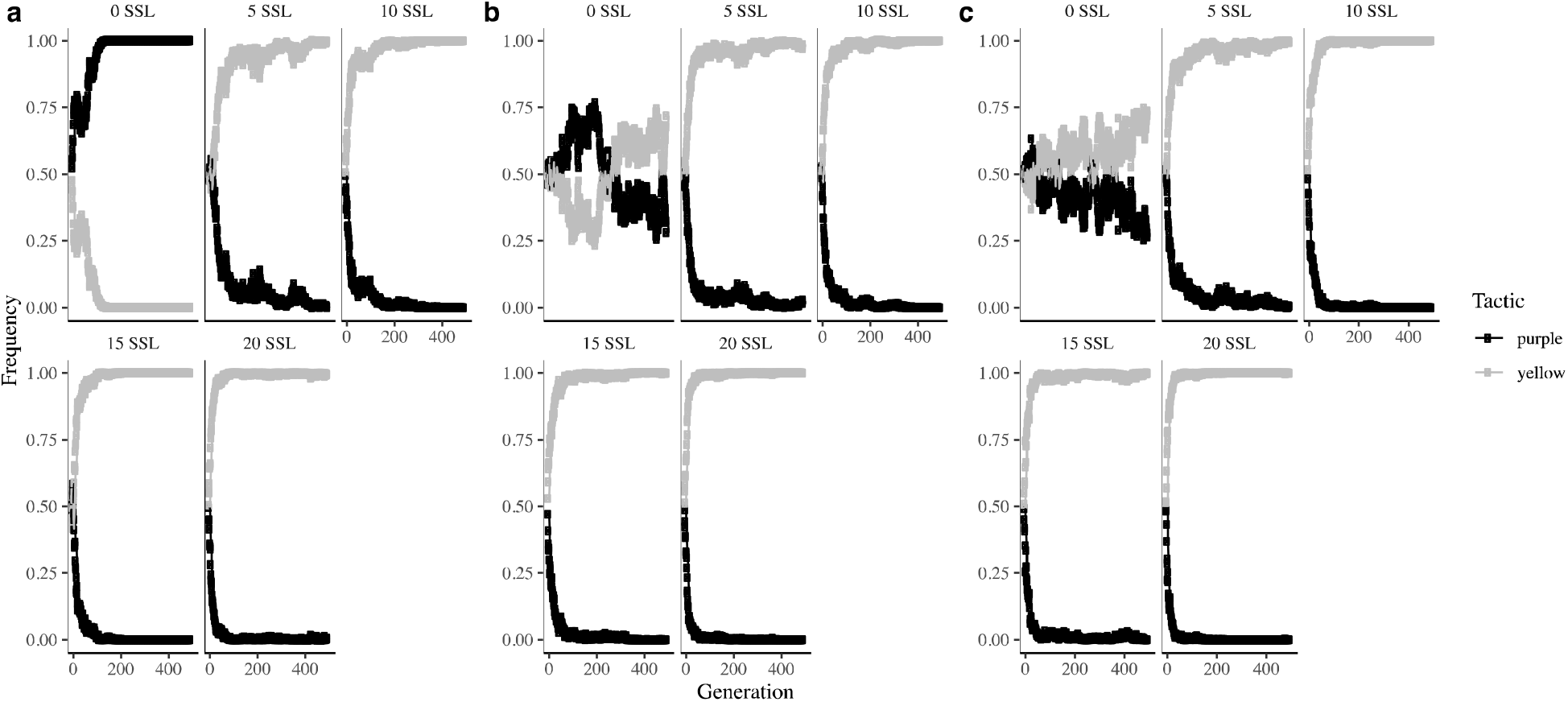

## Discussion

Alternative reproductive tactics abound in nature (Sinervo and Calsbeek 2010), yet few processes have been identified that sustain them. Those that have been identified – negative frequency-dependent selection and disruptive selection – both focus on selection acting directly on the sex exhibiting the polymorphism (Sinervo and Calsbeek 2006). We show that selection acting on both sexes, and intralocus sexual conflict that results when genetic architecture is shared, may also preserve sex-specific alternative reproductive tactics over hundreds of generations. If male ARTs differentially affect the fitness of their daughters, then they can be maintained (or eliminated) by selection for (against) their daughters even in the absence of negative frequency-dependent fitness. While this possibility has been touched upon in a few studies (Bielak et al. 2014; HeinenLJKay et al. 2020; Łukasiewicz et al. 2020), this is the first theoretical model to our knowledge that addresses the maintenance of ARTs through intralocus sexual conflict.

In addition to negative frequency-dependent selection, two other processes emerged from our model that preserved alternative tactics over 500 generations. First, both tactics persisted when there was no direct selection on males, and most (15) or all (20) loci were sex-specific in their expression. In this scenario, males experienced no correlated selection, because males could not pass on their liability trait value to their daughters. Fecundity selection favored females with high liability trait values, but these females could not pass on their liability trait values to their sons because intersexual heritability was zero. Thus, neither direct nor correlated selection acted on the liability trait in males, allowing the persistence of both tactics over 500 generations. The fact that purple males comprised the majority of males by the 500^th^ generation in most of these model runs suggests that these scenarios could not maintain ARTs indefinitely. However, because the trend toward an purple male majority occurred more slowly in these models as the number of sex-specific loci increased, low intersexual heritability could play a role in preserving male ARTs in natural populations where selection on males is chronically weak or temporally and spatially variable.

The second scenario in which both ARTs were maintained over 500 generations was when yellow males were 1.8 – 2 times more likely to survive and mate than purple males, and the number of sex-specific loci was zero. This created a scenario in which males experienced direct selection favoring yellow males and correlated selection favoring purple males. Because no loci were sex-specific in their expression, the liability trait was heritable between the sexes, creating intralocus sexual conflict. When the strength of directional selection on male tactic was lower (i.e., the fitness of yellow males was just 1.5 times higher than that of purple males), correlated selection outweighed direct selection and purple males went to fixation. However, as the strength of directional selection increased, direct and correlated selection balanced each other out, and both tactics persisted over 500 generations.

Persistent intralocus sexual conflict in these scenarios was evident in the maintenance of the sex load. Selection gradients on females remained positive and those on males remained negative throughout these simulations due to intralocus sexual conflict constraining the evolution of sexual dimorphism. In concordance with theory, the maintenance of the sex load in these simulations resulted in a lower population growth rate compared with simulations in which the sexes could respond independently to selection. This finding shows that the maintenance of ARTs, when achieved through intralocus conflict, may reduce population fitness.

Our model also showed that one male tactic could go to fixation even when males did not experience direct selection. When the liability trait was selected to be higher in females, but did not experience direct selection in males, purple males became fixed in the population when intersexual heritability for the liability trait was high. This occurred through correlated fecundity selection. This finding, though not explicitly predicted, again highlights the importance of investigating selection in both sexes when attempting to understand the evolutionary maintenance of sex-specific polymorphisms.

Our models consistently revealed that sexual conflict resulted in the persistence of a sex-specific polymorphism over hundreds of generations, even in the absence of negative frequency-dependent or disruptive selection. This result is important in light of a growing literature documenting genetic correlations between male ARTs and female fitness, and we suggest that these systems might be the most fruitful places to begin investigating the role of sexual conflict in ART maintenance. For example, side-blotched lizards (*Uta stansburiana*) exhibit three male morphs associated with orange, blue, and yellow throat colors (Sinervo and Lively 1996). In this system the intersexual heritability of throat color is high, and female throat color is correlated with fecundity, such that the genes associated with male ART have direct consequences for female fitness and population growth rate (Sinervo and Zamudio 2001). Though negative frequency-dependent selection seems partly responsible for ART maintenance in this system, genetic correlations between male tactic and female fitness also have the potential to affect ART frequencies (Sinervo and Zamudio 2001).

Male Pacific field crickets (*Teleogryllus oceanicus*) in Hawaii exhibit singing and silent (satellite) reproductive tactics that stem from different wing morphologies (normal and flatwing; Zuk et al. 2006). These wing morphologies have a genetic basis, and both males and females carry the genes associated with male wing morphology. Females carrying flatwing alleles invest less in reproductive tissue mass, experience more frequent mating failure, and are slightly less likely to mount and mate with males compared to normal females (Heinen-Kay et al. 2019a; HeinenLJKay et al. 2020; Richardson et al. 2021). The maintenance of both male ARTs in this scenario is likely due in part to sexual selection, as females prefer males that produce song (Tanner et al. 2019). Interestingly, negative frequency-dependent selection is unlikely in this system – while flatwing males must eavesdrop on calling males to find females (Zuk et al. 2006), both flatwing and normal males are less likely to employ satellite behavior when reared in an acoustic environment with frequent male calls, and more likely to employ satellite tactics when reared in silence (Bailey et al. 2010). Because flatwing males have much higher survival (Zuk et al. 2006) and sire more offspring than normal males (Heinen-Kay et al. 2019b), while females carrying flatwing alleles seem to have lower fitness, intralocus sexual conflict over wing morphology may also play a role in the maintenance of both male ARTs (HeinenLJKay et al. 2020; Richardson et al. 2021). However, we note that the survival benefit of the flatwing mutation in males likely outweighs the fitness detriment to females, and that sexual conflict over the flatwing allele is partially resolved due to demasculinized gene expression in flatwing females (Rayner et al. 2019) so this idea requires further investigation.

Studies of the bulb mite (*Rhizoglyphus* spp.) are also consistent with ART maintenance through intralocus sexual conflict. In artificial selection experiments, females from lines selected for fighter males have lower fitness (fecundity and longevity) than females from lines selected for sneakers (Bielak et al. 2014; Łukasiewicz et al. 2020). Harano et al. (2010) showed the same pattern in flour beetles (*Gnatocerus cornutus*) using an artificial selection experiment in which they selected for larger or smaller male mandibles. While flour beetles do not have ARTs *per se*, they do exhibit sexual dimorphism in a sexually selected weapon, and the intralocus sexual conflict arising from simultaneous sexually antagonistic selection and a negative intersexual genetic correlation between male mandible size and female fitness (via abdomen size and thus egg production) could maintain variation in male mandible size as shown in the present paper.

While the concepts of intralocus sexual conflict and correlated selection are decades old (Lande and Arnold 1983; Rice and Chippindale 2001), they have only just begun to penetrate the literature on alternative reproductive tactics. Our results show that intralocus sexual conflict can play a role in the maintenance of ARTs, and that understanding the maintenance – or elimination – of sex-specific polymorphisms requires considering selection in both sexes. Fruitful systems in which to look for this phenomenon in nature will include those in which the trait or traits that differentiate ARTs are also expressed in females (e.g., body size or condition), or those in which genetic correlations between ARTs and female survival or reproductive success have already been established.

## Data Availability Statement

The individual-based model used in this paper has been submitted with the manuscript and will be uploaded to Dryad upon acceptance.

